# Effects of Temperatures and High Pressures on the Growth and Survivability of Methanogens and Stable Carbon Isotope Fractionation: Implications for Deep Subsurface Life on Mars

**DOI:** 10.1101/281105

**Authors:** Navita Sinha, Sudip Nepal, Timothy Kral, Pradeep Kumar

**Author notes:** Correspondence and requests for materials should be addressed to P.K. [ ].

## Abstract

In order to examine the potential survivability of life in the Martian deep subsurface, we have investigated the effects of temperature (45°C, 55°C, and 65°C) and pressure (1 atm, 400 atm, 800 atm, and 1200 atm) on the growth, carbon isotopic data, and morphology of chemolithoautotrophic anaerobic methanogenic archaea, *Methanothermobacter wolfeii*. The growth and survivability of this methanogen were determined by measuring the methane concentration in headspace gas samples after the cells were returned to their conventional growth conditions. Interestingly, this methanogen survived at all the temperatures and pressures tested. *M. wolfeii* demonstrated the highest methane concentration following exposure to pressure of 800 atm and a temperature of 65°C. We found that the stable carbon isotopic fractionation of methane, δ^13^C(CH_4_), was slightly more enriched in ^12^C at 1 atm and 55°C than the carbon isotopic data obtained in other temperature and pressure conditions. A comparison of the images of the cells before and after the exposure to different temperatures and pressures did not show any obvious alteration in the morphology of *M. wolfeii*. The research reported here suggests that at least one methanogen, *M. wolfeii*, may be able to survive under hypothetical Martian subsurface conditions with respect to temperature and pressure.

## Introduction

The current surface conditions of Mars are extremely harsh for any known life forms. Environmental factors such as low surface temperature, low atmospheric pressure, DNA damaging UV radiation, and the presence of oxidizing compounds make Mars an inhospitable planet (Biemann and others 1977; Cockell and others 2000; Jakosky 1998). If life exists on the surface of Mars, it would have to challenge the environmental extremes of Mars and would have unique adaptation mechanisms. Therefore, the most feasible possibility of finding active known life forms on Mars would be “near or deep subsurface”, where the temperature and pressure will be higher compared to the surface and the environment will be protected from the damaging cosmic radiation. The discovery of extremophiles and the knowledge of Earth’s subsurface biospheres have also bolstered the idea of searching for life in the subsurface of other planetary bodies such as Mars (Cavicchioli 2002).

The detection of methane in the Martian atmosphere (Formisano and others 2004; Krasnopolsky and others 2004; Mumma and others 2004; Mumma and others 2009; Webster and others 2015) has further reinforced the search for extinct or extant life on Mars. The reason is that most terrestrial methane is produced by biological sources either directly or indirectly (Atreya and others 2007). Methanogens are one of the various sources of methane on Earth. Some strains of methanogens have shown survivability and growth in simulated Martian physical and chemical conditions (Kral and Altheide 2013; Kral and others 2004; Kral and others 2014; Kral and others 2015; McAllister and Kral 2006; Sinha and Kral 2015). For these reasons, methanogens, which are chemolithoautrophic anaerobic archaea, have been considered ideal candidates for life on Mars (Boston and others 1992; Chapelle and others 2002; Chastain and Kral 2012; Kral and others 2004; Kral and others 2015; Moran and others 2005). The sources of methane on Mars are still unknown. Mars’ atmospheric methane could be the results of biotic, abiotic or a combination of both processes. Methanogens could be one of the several potential sources of Martian methane.

Stable carbon isotope fractionation is one of the several potential techniques to differentiate between biogenic and abiogenic sources of methane (Allen and others 2006). Stable carbon isotope fractionation data for terrestrial atmospheric methane have been used in order to understand the environments, pathways, and origins or substrates of methanogenesis (Londry and others 2008; Rothschild and DesMarais 1989; Schidlowski 1992). Sinha and Kral have recently studied the carbon isotope fractionation following methanogenesis on various Mars regolith analogs and found enriched values of ^12^C on the clay called montmorillonite compared to the carbon isotopic data obtained on other Mars analogs such as JSC Mars-1, Artificial Mars Simulant, and Mojave Mars Simulant (Sinha and Kral 2015).

Analogous to Earth’s subsurface environments, hydrothermal systems might have existed and may also be present on Mars. On Earth, several hyperthermophilic and barophilic archaea have been isolated and characterized from the deep sea floor and hydrothermal sites (Canganella and others 1997; Horikoshi 1998; Shimizu and others 2011; Takai and others 2002). Some of the most abundant species near hydrothermal vents are hyperthermophilic methanogens (Takai and others 2004). Life near a hydrothermal vent experiences a wide range of temperature and pressure. It has been found that a methanogen of genus *Methanopyrus* can grow up to a temperature of 110°C (Takai and others 2008) and *Methanococcus jannaschii* demonstrated methanogenesis up to a pressure of 750 atm (Miller and others 1988). Methanogenesis has also been found in organisms thriving in 3.5-million-year-old subseafloor basalt on Earth and was detected by using δ^13^C data (Lever and others 2013). Several surface features, geochemical, and isotopic evidence in a Martian meteorite point to the activity of hydrothermal systems on Mars (Brakenridge and others 1985; Romanek and others 1994; Shock 1997; Watson and others 1994). Therefore, similar subsurface biota might exist in the Mars’ subsurface.

There are a few studies done on the effect of temperature and pressure on various strains of methanogens. However, the effects of a wide range of temperature and high pressure on methanogens in the context of Mars’ subsurface have not been studied before. The goal of this study is to examine the growth and survivability of a Mars’ model organism, *Methanothermobacter wolfeii* in a wide range of temperatures (45°C - 65°C) and pressures (1 atm, - 1200 atm). *M. wolfeii*, a hydrogenotrophic methanogenic archaeon used in this study utilizes CO_2_ for its carbon source, H_2_ for its energy source and produces methane as a metabolic byproduct. We have also measured the stable carbon isotope fractionation of methane, δ^13^C(CH_4_), in order to understand the effect of optimal and non-optimal temperature and pressure on the carbon isotopic data. The images of cells were also acquired and were analyzed to investigate morphological changes following exposure to various pressures and temperatures.

## Materials and Methods

### Preparation of a stock culture of *M. wolfeii*

*M. wolfeii* (OCM36) was obtained from the Oregon Collection of Methanogens, Portland State University, Portland, OR and was grown in MM medium consisting of potassium phosphate, ammonium chloride, calcium chloride, resazurin as an oxygen indicator, and many trace minerals (Xun and others 1988) in a bicarbonate buffer. MM medium was prepared in an anaerobic chamber (Coy Laboratory Products Inc., Grass Lake Charter Township, MI), which was filled with 90% carbon dioxide and 10% hydrogen. The growth medium was then transferred into anaerobic culture tubes inside the anaerobic chamber as described previously ((Boone and others 1989). The tubes were then sealed with butyl rubber stoppers, removed from the chamber, crimped with aluminum caps, and autoclaved for sterilization.

A sterile sodium sulfide solution (2.5% wt/vol; 0.15 mL per 10 mL media) was then added to each tube about an hour prior to inoculation of the methanogen (Boone and others 1989) in order to eliminate any residual molecular oxygen from the tubes containing the media. After inoculating the methanogen, the tubes were pressurized with 2 atm of H_2_ gas and incubated at the organism’s optimal growth temperature, 55°C. The stock culture of *M. wolfeii* was maintained by transferring into fresh MM medium every fifteen days.

The growth and survivability of this methanogen were determined by measuring the methane concentration in an aliquot of the headspace gas using a gas chromatograph (Shimadzu 2014, USA), which was equipped with a flame ionization detector.

### Temperature-Pressure experiments

The schematic diagram of the temperature-pressure chamber is shown in Figure 1. A quartz cuvette (Spectracell, Oreland, PA) was filled with 1 mL of fresh liquid culture of *M. wolfeii* and was capped with a Teflon cap (E. I. DuPont de Nemours, Paris, France) inside the anaerobic chamber. The cuvette filled with sample was then put into a high hydrostatic pressure-temperature chamber (ISS, Champaign, IL), which was filled with water and was connected to a piston. The pressure inside the chamber was developed with the help of a piston by pressurizing liquid water, and the pressure was measured with a pressure gauge attached to the piston (Kumar and Libchaber 2013). The temperature of the chamber was maintained using a circulating water bath (Neslab, USA) and was measured in real time using a thermocouple (National Instruments, USA) attached to the chamber. Before loading the sample, the temperature of the chamber was equilibrated to the desired temperature and the pressure was applied after the sample cuvette was put into the high temperature-pressure chamber. The system equilibrated to the desired pressure and temperature in less than two minutes after loading the sample. In this work, we preformed experiments at three different temperatures—45°C, 55°C, and 65°C and four different pressures—1 atm, 400 atm, 800 atm, and 1200 atm; resulting in a total of twelve sets of experiments. For a given temperature and pressure, the sample was kept in the temperature-pressure chamber for 15 hours. After this time, the pressure of the chamber was released and the sample-filled cuvette was removed from the chamber.

**Figure 1.**
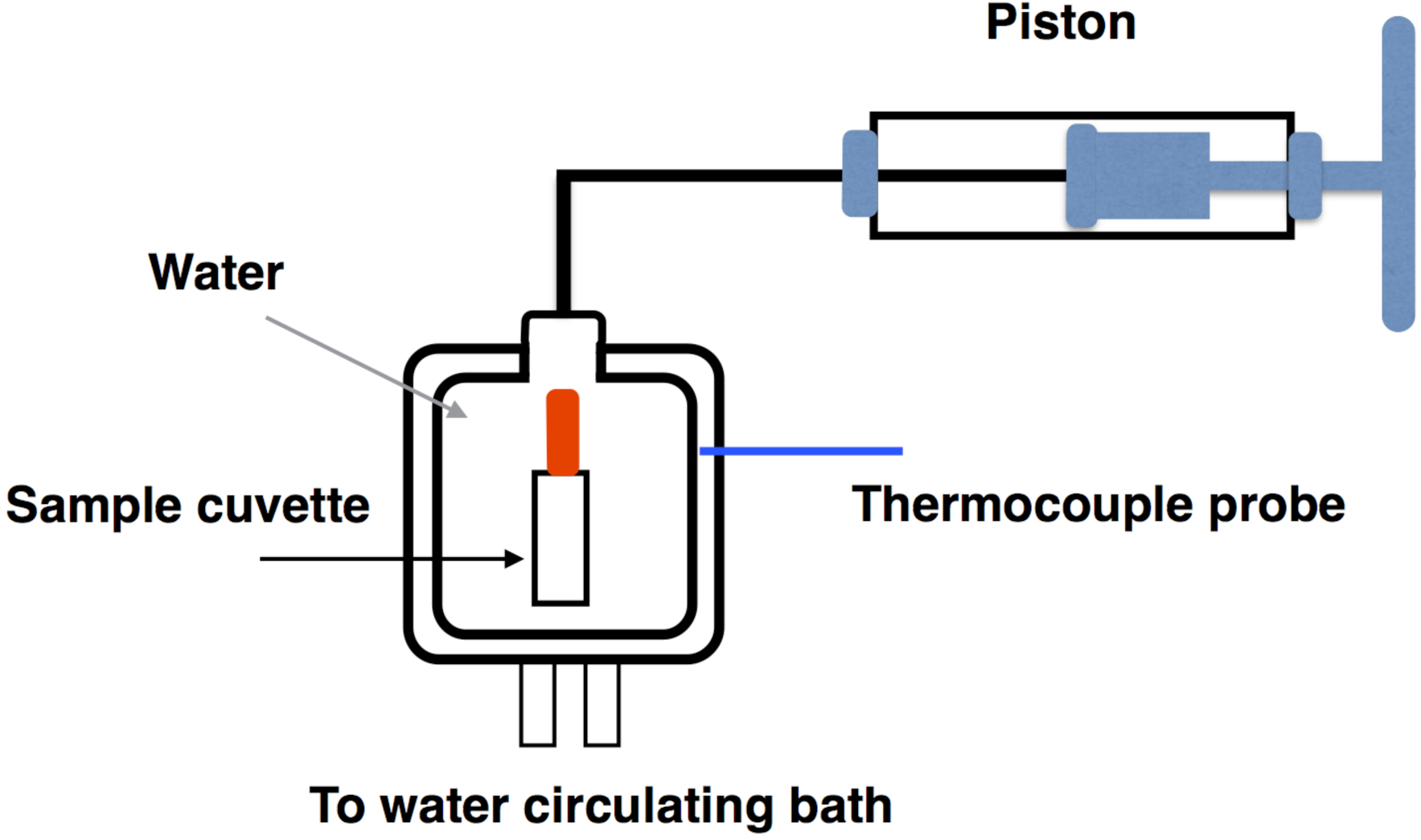
Experimental setup for the temperature-pressure experiment.

The cells from the cuvette were then transferred to fresh medium. Five hundred microliters of cells from the cuvette were mixed with 500 uL of sterilized MM medium in a vial to make a total of 1 mL of culture. From this, 300 uL cells were inoculated into three different anaerobic tubes containing 10 mL of sterilized MM medium to make triplicate samples. The tubes were then pressurized with 2 atm H_2_ gas and incubated at the conventional growth temperature, 55°C. The growth and survivability were determined by measuring methane concentration in the headspace gas of each sample at regular intervals with the help of a gas chromatograph (Shimadzu 2014).

### Determination of stable carbon isotope fractionation

Using the procedure described by Sinha and Kral (2015), the carbon isotope fractionation of methane in the headspace gas of all samples was measured periodically by a Piccaro Cavity Ringdown Spectrometer G-2201-I isotopic CO_2_/CH_4_ in the University of Arkansas Isotope lab. The carbon isotope fractionation, δ^13^C, was calculated using the following equation:

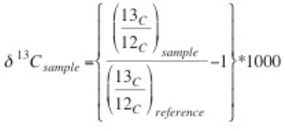

The reference isotopic standard for δ^13^C is Pee Dee Belemonite (O’Leary 1981).

### Imaging of cells

Phase contrast images of the cells were obtained before and after exposure to various temperatures and pressures using a SPOT Imaging camera and 40X objective mounted on a Nikon Optiphot microscope.

## Results

*M. wolfeii* was exposed to temperatures of 45°C, 55°C, and 65°C and pressures of 1 atm, 400 atm, 800 atm, and 1200 atm in a high hydrostatic pressure-temperature chamber for 15 hours. Interestingly, *M. wolfeii* survived at all temperatures and pressures studied here. This methanogenic archaeon demonstrated methanogenesis by producing methane after the cells were returned to their conventional growth conditions. All measurements of methane concentration were taken post exposure to different temperatures and pressures. In Figure 2, we show methane concentration, [CH_4_], as a function of time for the cells exposed to the temperatures—45°C, 55°C, and 65°C. Data in these figures represent the average methane concentration produced by methanogens exposed to the pressures—1 atm, 400 atm, 800 atm, and 1200 atm. For all temperatures and pressures, samples were in triplicates. For each temperature, *M. wolfeii* demonstrated highest methane concentration following the exposure to 800 atm of pressure and the lowest methane concentration following 1 atm. Highest methane concentration varies for different temperatures and reaches the maximum on different days. For example, at 800 atm and 45°C and 55°C, methane concentration reached its maximum on the third day while for 65°C the methane concentration reached its maximum on the fourth day. Moreover, we found that at 55°C, the optimal growth temperature of *M. wolfeii*, the lag phase was little more than 24 hours whereas at 45°C and 65°C (non-optimal growth temperatures), the lag phases were less than 24 hours. This suggests that after the release of non-optimal physical conditions (stress), *M. wolfeii* adapted quickly to the optimal growth condition.

**Figure 2:**
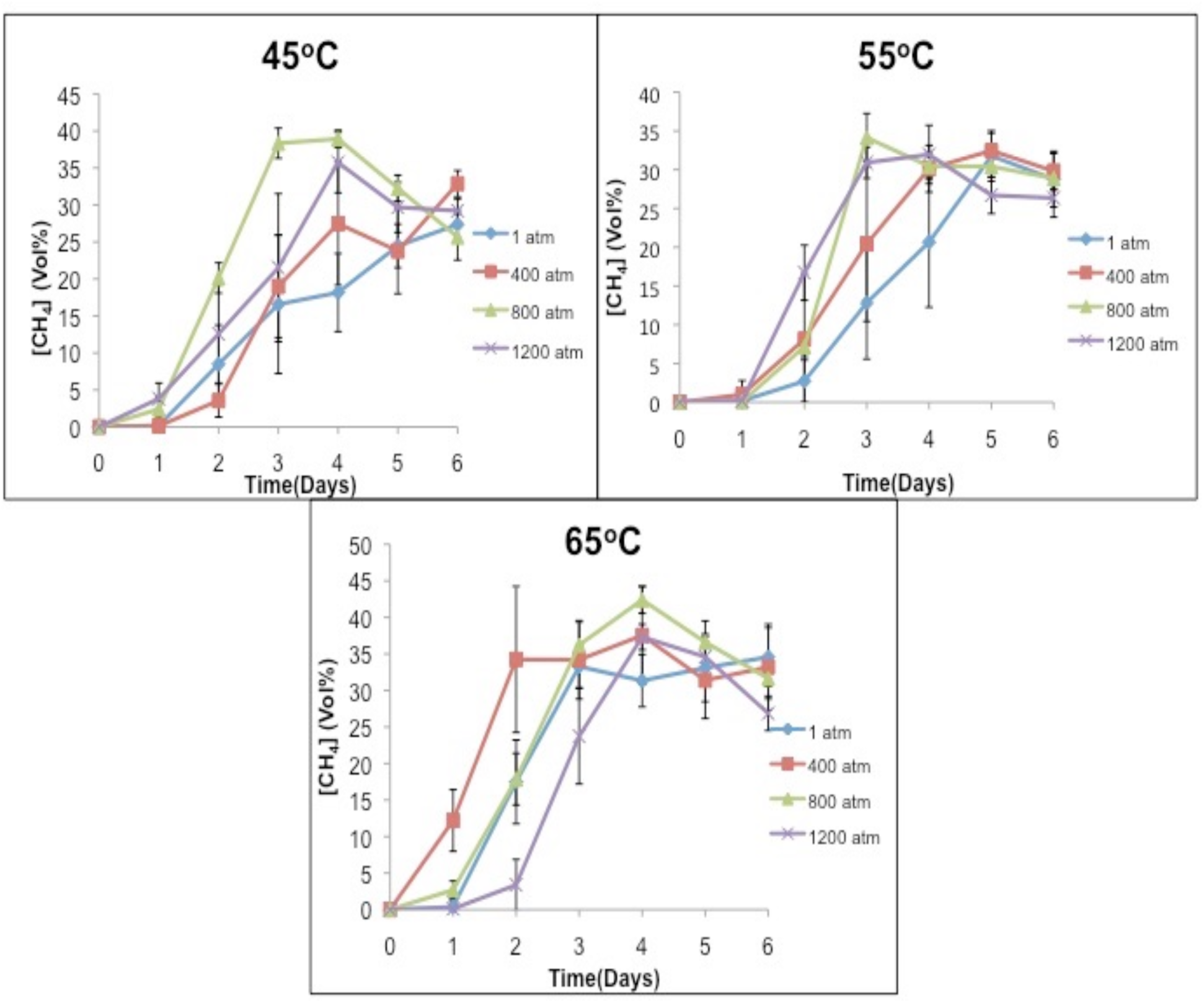
Methane concentration as a function of time following methanogenesis for *Methanothermobacter wolfeii* exposed to the temperatures—45°C, 55°C, and 65°C. The different colored lines in these figures represent the average methane concentration produced by methanogens exposed to the pressures—1 atm, 400 atm, 800 atm, and 1200 atm.

In Figure 3, we show methane concentration, [CH_4_], as a function of time for the cells exposed to the pressures—1 atm, 400 atm, 800 atm, and 1200 atm. Data in these figures represent methane produced by methanogens exposed to the temperatures—45°C, 55°C, and 65°C. For all temperatures and pressures, samples were in triplicates. For each pressure, *M. wolfeii* exhibited the highest methane concentration after exposure to 65°C. For the lowest pressure, 1 atm, the methane concentration reached its maximum on the third day following a temperature of 65°C. However, for the highest pressure, 1200 atm, the methane concentration reached its maximum on the fourth day for all of the temperatures tested.

**Figure 3:**
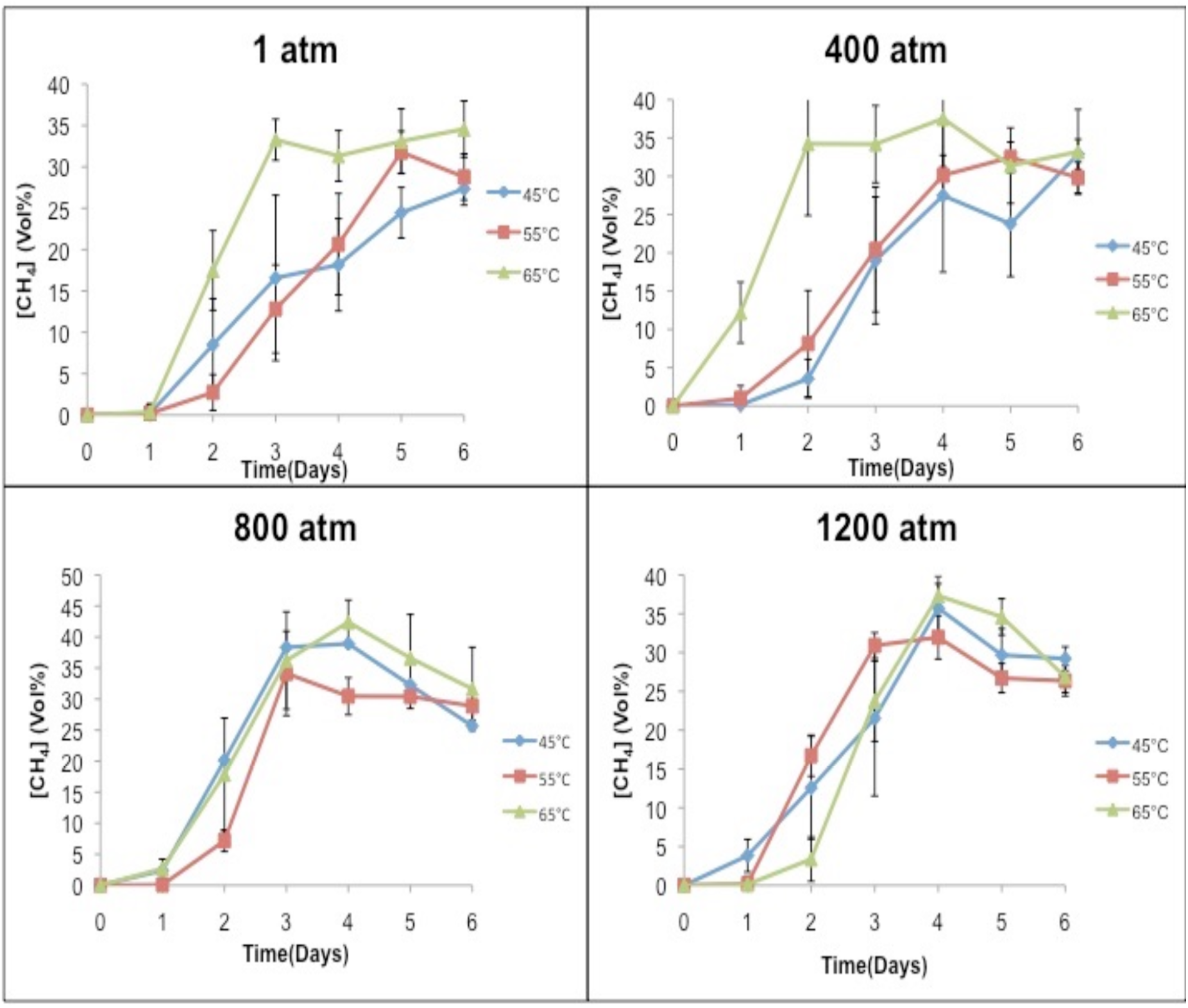
Methane concentration as a function of time following methanogenesis for *Methanothermobacter wolfeii* exposed to the pressures—1 atm, 400 atm, 800 atm, and 1200 atm. The different colored lines in these figures represent the average methane concentration produced by methanogens exposed to the temperatures—45°C, 55°C, and 65°C.

We next studied the effect of temperature and pressure on the stable carbon isotope fractionation of methane produced by *M. wolfeii*. In Table 1, we have listed the carbon isotope fractionation data obtained on the second and sixth day following the pressure-temperature exposures. For 1 atm and 55 °C (conventional growth conditions of *M. wolfeii*), δ^13^C(CH_4_) on the second and sixth day are -74.36‰ and -73.35‰, which are comparable to published data for standard growth conditions (Sinha and Kral 2015). Here we found slightly depleted values of δ^13^C(CH_4_) in the conventional conditions compared to the δ^13^C(CH_4_) data obtained at non-optimal temperatures and elevated pressures.

**Table 1:** Carbon isotope fractionation of methane, δ^13^C(CH_4_), produced by *Methanothermobacter wolfeii* obtained on Day 2 and Day 6 following different temperature-pressure exposures. Values shown are in per mil.

|  | **45°C** |  | **55°C** |  | **65°C** |  |
| --- | --- | --- | --- | --- | --- | --- |
| Pressure (atm) | Day 2 | Day 6 | Day 2 | Day 6 | Day 2 | Day 6 |
| 1 | -70.86 | -70.74 | -74.36 | -73.35 | -70.43 | -70.94 |
| 400 | -72.30 | -71.98 | -70.96 | -71.04 | -69.38 | -69.66 |
| 800 | -72.88 | -73.06 | -73.25 | -71.55 | -71.42 | -69.72 |
| 1200 | -73.74 | -69.79 | -69.13 | -70.64 | -69.55 | -69.48 |

In order to compare any morphological changes in the cells exposed to optimal and non-optimal pressures and temperature, we obtained phase contrast images of the cells before and after exposure to different temperature and pressure. In Figure 4, we compare the images of cells exposed to temperatures—55°C and 65°C, and pressures—1 atm and 800 atm. We did not find any significant effect of temperature and pressure on the morphology of *M. wolfeii.* It has been shown that high pressure exhibits cell division inhibiting effects on mesophilic bacteria such as E. coli (Kumar and Libchaber 2013). The lack of morphological alteration such as elongation in the case of M. *wolfeii* suggests that the exposure time of 15 hours was not long enough for the cells to undergo multiple cycles of cell division.

**Figure 4:**
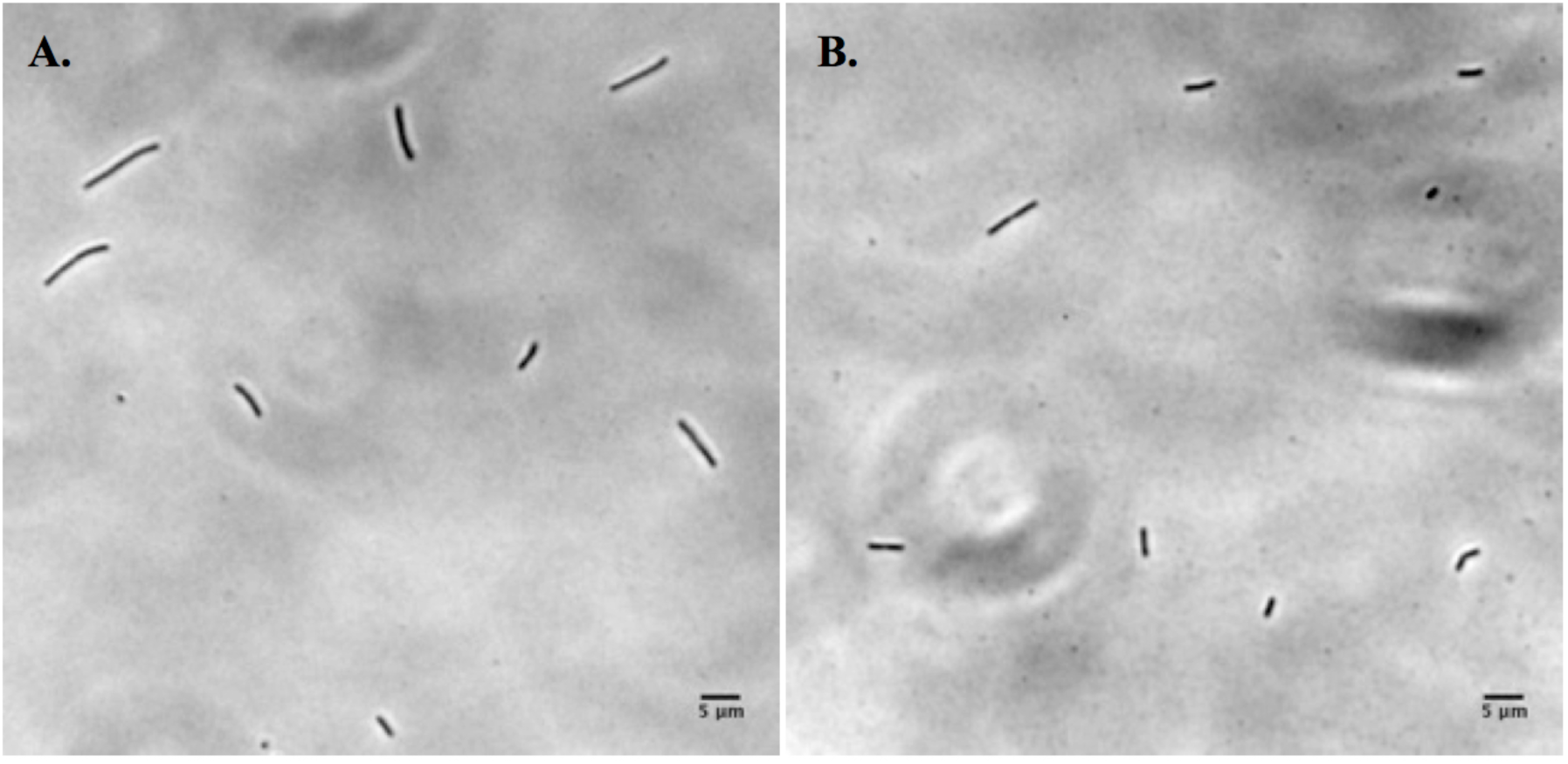
Images of *Methanothermobacter wolfeii* after exposure to (A) T=55°C and P=1 atm (B) T=65°C and P=1200 atm under a magnification of 400X.

## Discussion

The growth and survivability of several methanogenic archaea in simulated Martian surface conditions such as at low temperature, low pressure, and desiccation have been studied previously (Kendrick and Kral 2006; Kral and others 2011; Kral and others 1998; Reid and others 2006). On the other hand deep subsurface of Mars could potentially offer a feasible environment for a biosphere. The challenges are then high pressure, temperature, and availability of liquid water, nutrients, and the source of energy. It is imperative to locate liquid water in the subsurface of Mars. All terrestrial life needs water at some stage in their life cycle. Martian geophysical models suggest that the liquid water in the subsurface of Mars could be present to a depth of ∼310 km (Jones and others 2011). According to this model, the depth and pressure for approximate temperature range 45°C-65°C would be between 1-30 km and 100-3,000 atm.

In this work, we have investigated the effects of temperature (45°C, 55°C, and 65°C) and pressure (1 atm, 400 atm, 800 atm, and 1200 atm) on the growth, carbon isotopic data, and morphology of *M. wolfeii*. The growth and survivability of methanogens were determined by measuring methane concentration in the headspace gas samples after they were returned to their conventional growth conditions. Due to the limitation on the maximum temperature and prolonged experiments in hydrostatic temperature-pressure chamber, 15 hours long experiments were performed with a maximum temperature of 65°C and pressure of 1200atm. Since the typical doubling time for *M. Wolfeii* is about 10 hours at conventional growth conditions, our experiments should able to detect the survivability over the time scale of our experiments. We found *M. wolfeii* was able to endure a temperature of 65°C and a pressure of 1200 atm for at least that amount of time. Surprisingly, *M. wolfeii* demonstrated methanogenesis following exposure to all the temperatures and pressures studied here. *M. wolfeii* exhibited the highest methane concentration following exposure to a pressure of 800 atm and a temperature of 65°C. The exact reason for this is not clear and will be the focus of future studies.

The stable carbon isotope fractionation of methane was measured in different temperature and pressure experiments. We found very slight or negligible difference in the carbon isotopic data following optimal and non-optimal growth conditions of *M. wolfeii.* The δ^13^C(CH_4_) data in the optimal conditions were slightly lower than the δ^13^C(CH_4_) data in non-optimal conditions. It is possible that for *M. wolfeii*, 15 hours exposure to various temperatures and pressures may not be long enough to have a significant effect on the carbon isotopic data. (Takai and others 2008) have found that *Methanopyrus kandleri* produced isotopically heavier methane under high hydrostatic pressure conditions compared to the methane produced by *M. kandleri* in a conventional growth condition (Takai and others 2008).

We compared the morphology of cells before and after the exposure to different temperatures and pressures. Most of the cells remained intact and we did not find obvious alteration in the morphology of M. *wolfeii*, unlike *E. coli*, which shows increase in cell length with increase in pressure (Kumar and Libchaber 2013). Since the doubling time of *E. coli* is very small as compared to *M. wolfeii*, it was possible to detect stochasticity in cell division and elongation of *E. coli* at high pressure over a large number of generation times.

The results presented here suggest that one Mars’ model microorganism, *M. wolfeii*, can survive under presumed Martian subsurface conditions in terms of temperature and pressure. Therefore, the search for life on Mars should also be focused on the deep-subsurface of Mars.

## Conclusions

Methanogens have been considered ideal life forms on Mars for a long time. Here, we have examined the growth and the survivability of a methanogen, *M. wolfeii*, in presumed deep-subsurface environments in terms of temperature and pressure. We used three different temperatures (45°C, 55°C, and 65°C) and four different pressures (1 atm, 400 atm, 800 atm, and 1200 atm). *M. wolfeii* demonstrated survivability by producing methane following exposure to all different temperatures and pressures. The growth and survivability of *M. wolfeii* were investigated after returning them to their conventional growth conditions. We have also measured the carbon isotopic fractionation of methane produced by *M. wolfeii* and found that δ^13^C(CH_4_) in optimal growth conditions was slightly lower than the values obtained in non-optimal growth conditions. A comparison of the images of cells before and after the exposure to different temperatures and pressures did not reveal any apparent alteration in the morphology of *M. wolfeii*.

## Acknowledgments

The authors would like to thank Erik Pollock, manager of the University of Arkansas Stable Isotope Laboratory, for his assistance in obtaining isotope data.

